# A cross-species protocol for ultrasound-guided intrauterine injections across gestation

**DOI:** 10.64898/2026.07.07.737050

**Authors:** Ana Rita Ribeiro Gomes, Natalie Hamel, Surjeet Mastwal, David C. Ide, Kuan Hong Wang, David A. Leopold

## Abstract

This step-by-step protocol provides a cross-species, non-surgical approach that enables prenatal gene delivery to the developing nervous system in rats and marmosets. Under transabdominal ultrasound guidance, intracerebroventricular viral injection (FIVI) of recombinant adeno-associated virus vectors into the fetal brain achieves robust and long-term transduction from prenatal stages into adulthood. This approach can be adapted to multiple species and target sites outside nervous system, enabling safe and selective intrauterine manipulation and the generation of diverse experimental models for basic and preclinical research.

For complete details on the use and execution of this protocol, please refer to Ribeiro Gomes et al (2026)^1^.

**Graphical abstract:** 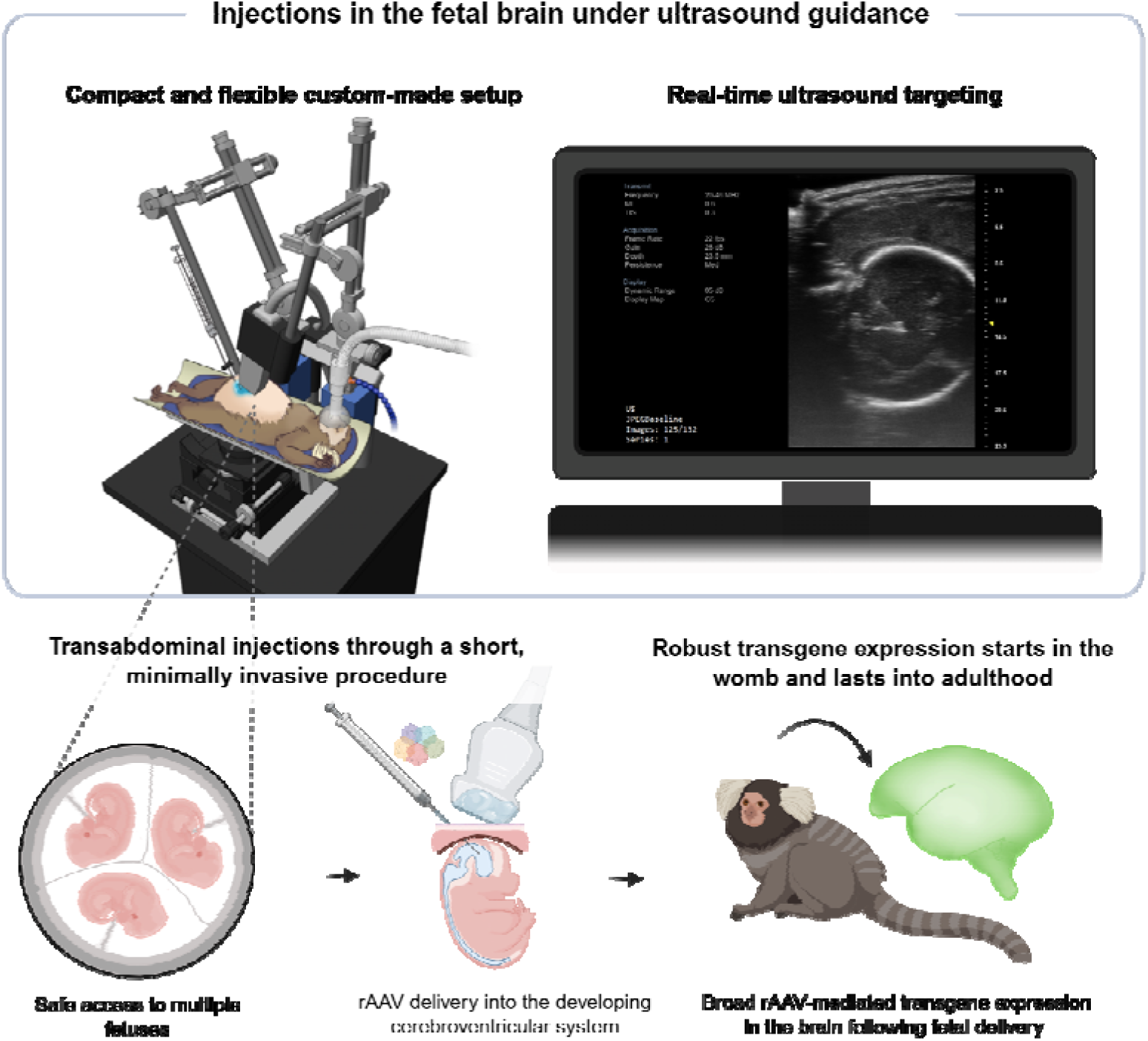

## Before you begin

Experimental procedures during gestation allow researchers to study developmental processes, including how manipulations of the fetus and its intrauterine environment influence biological outcomes. Ultrasound imaging guidance greatly facilitates such interventions by providing safe and targeted access to fetal compartments, including for prenatal gene delivery to developing neural cell populations. Critically, delivery of recombinant adeno-associated viruses (rAAVs) into the cerebrospinal fluid (CSF) of developing animals enables widespread gene transfer across the brain. The efficiency and distribution of transduction are strongly influenced by developmental stage, making the timing of delivery an important experimental variable. In altricial species such as mice, major developmental processes, including cortical lamination and the establishment of long-range connections, begin prenatally but continue throughout early postnatal life. In primates, however, development is more advanced at birth, and many equivalent developmental events are shifted to the prenatal period. Consequently, develop ental stages that can be targeted postnatally in mice require prenatal access in primates. Here, we present a step-by-step protocol for ultrasound-guided intracerebroventricular viral injection (FIVI) of rAAV in fetal marmosets (Callithrix jacchus) and rats (Rattus norvegicus). The procedure was initially developed and optimized in rats before being translated to marmosets, small New World primates that share key developmental, anatomical, and functional characteristics with humans. Together, these models illustrate the cross-species applicability of the approach, while providing gene delivery strategies for both a genetically tractable rodent model and a translationally relevant nonhuman primate. FIVI enables broad gene transfer and stable, long-term transgene expression in wild type animals, facilitating the generation of complementary quasi-transgenic models for research and translational applications from prenatal development through adulthood.

### Innovation

The key innovation of this protocol is the ability to perform fetal injections in a nonhuman primate model through a short, minimally invasive, non-surgical procedure, addressing the longstanding challenge of safe and targeted access during gestation. A custom mechanical setup stabilizes the injection trajectory, while electrocautery-assisted entry facilitates penetration of maternal tissues with minimal displacement of the fetus. Combined with ultrasound guidance, this approach enables selective targeting of fetal structures while avoiding laparotomy, uterotomy, and direct manipulation of the uterus or fetuses. Open surgical approaches provide access to the fetus but increase procedural complexity and require direct manipulation of the uterus and/or fetus^2-4^. In nonhuman primates, these constraints have historically limited the adoption of fetal manipulation approaches, which remain rarely performed and have consequently yielded few experimental models for investigating developmental processes relevant to human biology and disease.

As an additional feature, although this protocol focuses on CSF injections, the approach can be adapted to target tissues beyond the fetal nervous system, including other fetal organs and components of the intrauterine environment such as the amniotic cavity and placenta. Initially developed in rats and subsequently adapted to marmosets, it establishes a cross-species framework for ultrasound-guided intrauterine manipulation and gene delivery that can be extended to additional species, thereby expanding the pool of genetically engineered models available for research.

Here, we describe the overall procedure and specify the steps required for fetal CSF injections in the two models used, marmosets and rats. The same core materials and equipment were used in both species, including the preparation and FIVI setups, ultrasound system and transducers, and guide and injection needles, with the same overall injection strategy unless otherwise noted. Required species-specific modifications are indicated at the corresponding steps throughout the protocol.

### Institutional permissions

All procedures were approved by the Animal Care and Use Committee (ACUC) of the Intramural Research Program of the National Institute of Mental Health (NIMH) and performed at the U.S. National Institutes of Health in accordance with institutional guidelines.

### Animals

Timed-mated Sprague-Dawley rats (Strain Code: 001) were purchased from Charles River Inc with gestational ages varying between postconception (PC) day 3 and PC12. For in-house breeding, 2-month male and female Sprague-Dawley rats were purchased from Charles River Inc. Dams used for FIVI procedures were 2-9 months old and weighted 300-500 grams. Upon arrival, rats were single-housed and acclimated to the vivarium for at least 72h before FIVI or time-mating procedures. Rats were housed in a 12 h light/12 h dark cycle and had ad libitum access to food and water.

Common marmosets (Callithrix jacchus) 2-9 years old from the NIMH intramural colony weighing 360-620 grams at the time of the procedures were used in the experiments. Marmosets were housed in a 12 h light/12 h dark cycle and had ad libitum access to food and water.

### In-house rat breeding

#### Timing: Prior to procedure

1. Synchronize 2–12-month-old female by intraperitoneal injection of 0.2 mL of a 0.4 mg/mL solution of [des-Gly10, D-Ala6]-LH-RH ethylamide acetate salt hydrate between 10:30 AM and 11:00 AM.

2. At 96 h after hormone administration, pair each female with a male between between 11:30 AM and 12:00 P.M.

**Note:** Breeding efficiency improves when males are single-housed and rested from mating for at least 1 week before pairing.

3. Check for a vaginal plug the following morning before 9:00 AM. If a vaginal plug is detected, designate that day as PC1, assume pregnancy, and schedule the FIVI procedure according to the desired gestational stage.

***Note.*** Vaginal plugs may not always be detected. If no vaginal plug is observed, pregnancy cannot be excluded. Weight the female every two days, and if body weight progressively increases, an ultrasound scan is performed under anesthesia on the date of the planned procedure to confirm pregnancy. For scheduling purposes, designate the morning on which the vaginal plug would have been detected as PC1 even when no plug is observed. Rats are not habituated to ultrasound exams. Therefore, pregnancy is not routinely confirmed by ultrasound before the planned procedure to avoid additional anesthesia or restraint of untrained pregnant animals.

### Marmoset ultrasound pregnancy checks

#### Timing: Prior to procedure

If marmosets are continuously paired for breeding rather than time-mated, use transabdominal ultrasonography to detect pregnancy and estimate gestational age. In our colony, breeding females are routinely examined by ultrasound at least once monthly regardless of known pregnancy status.

1. Transfer the female from the housing room to a procedure room containing the ultrasound system.

2. Gently hand-restrain the female while minimizing movement. Provide a positive food reward (e.g., dried fruit or Pediasure) before, during, and after the examination.

3. Perform transabdominal ultrasonography using the FUJIFILM VisualSonics Vevo MD system with a UHF22 or UHF48 linear transducer.

4. Measure the ventro-dorsal and transverse diameter of the uterus and uterine lumen, and, when visible, the biparietal diameter of the fetal skull. Use these measurements to estimate gestational age.

5. Estimate PC day by comparing the measured values with published measurements from Oerke et al. (1995)^5^.

**Note:** Based on established ultrasound measurements from Oerke et al. (1995)^5^, pregnancy can be detected as early as approximately PC14 and is typically readily apparent by PC21 based on the presence of an open uterine lumen. The PC day at the time of the procedures is estimated using these values. Subsequently, these dates are re-estimated after birth by subtracting 143 days from birth, accounting for the average pregnancy length in marmosets, between 140 and 145 days^5^.

### Preparation of the pregnant dam

#### Timing: Prior to procedure

1. Female marmosets are continuously paired with males for breeding in stable family groups and are not hormonally synchronized. Timed-mated rats are either purchased from an approved vendor or bred in-house.

### Viral vector selection and storage

#### Timing: Prior to procedure

The viral vector should be selected according to the experimental application. The rAAV vectors listed in the Key Resources Table were used to generate the representative outcomes shown in this protocol but is not required for performing FIVI.

2. Obtain a purified, high-titer viral vector encoding the gene of interest from a commercial supplier or viral vector core. Upon receipt, the viral vector encoding the gene of interest is stored at -80°C until use.

**Note.** The choice of rAAV construct, including serotype, promoter, transgene, and viral titer, should be determined according to the experimental application. Similarly, the gestational stage for injection should be selected according to the experimental objective, vector used, and cell populations intended for transduction. New vectors or applications may require empirical optimization of injection timing. The viral vectors used to validate this protocol were obtained as pre-prepared viral vector preparations. rAAV production and purification are therefore outside the scope of this protocol.

### Preparation of sterile FIVI instruments

#### Timing: Prior to procedure

3. Sterilize all procedural instruments – needles, needle guides, syringes, filtered pipette tips, curved hemostatic forceps, ball end hex driver 5/64" and # 3/32 – using an autoclave or ethylene oxide gas sterilization.

4. Verify that all procedural drugs (see Key Resources Table) and consumables – sterile and non-sterile tongue depressors, sterile and non-sterile cotton swabs, gauze, sterile and non-sterile ultrasound gel, disinfectant wipes, return electrode – are available and within their expiration dates.

5. Prepare the setup equipment and disinfect external surfaces by wiping with sodium hypochlorite-based disinfectant wipes.

**Note.** The disinfectant used should be selected in accordance with the institutional guidelines and facility-specific standard operating procedures.

## Key resources table

| REAGENT or RESOURCE | SOURCE | IDENTIFIER |
|---|---|---|
| **Bacterial and virus strains** |  |  |
| AAV2-CAG-GFP | Addgene | 37825-AAV2 |
| **Chemicals, peptides, and recombinant proteins** |  |  |
| Atropine Sulfate Injection, 0.54 mg/mL | MWI | NDC 13985-354-04 |
| Meloxicam Injection USP, 5 mg/mL | Meloxivet | NDC 17033-051-10 |
| Diazepam Injection, 5 mg/mL | Multiple sources | N/A |
| 50% dextrose | Multiple sources | N/A |
| 0.9% sodium chloride solution | Multiple sources | N/A |
| Isoflurane | Fluriso | NDC 13985-528-40 |
| Hair removal creme (Nair) | Church & Dwight Co., Inc. | N/A |
| Ophthalmic Ointment (0.4% Sodium Hyaluronate) | Ophthavet | NDC 17033-047-05 |
| Ethanol wipes | Multiple sources | N/A |
| Povidone iodine | Purdue Products LP | NDC 67618-154-16 |
| Lidocaine Ointment USP, 5% | Multiple sources | N/A |
| Water-based lubricant | Multiple sources | N/A |
| Non-sterile ultrasound gel | Multiple sources | N/A |
| Aquasonic 100 - Sterile Single Use - Overwrapped Foil Pouches 48 per box | Aquasonic | PLI 01-01 |
| 10 x PBS | Multiple sources | N/A |
| **Experimental models: Organisms/strains** |  |  |
| Sprague-Dawley rats | Charles River | Strain Code: 001 |
| Common marmosets (Callithrix jacchus) | NIMH | N/A |
| **Software and algorithms** |  |  |
| Fiji | ImageJ | https://imagej.net/software/fiji/ |
| Adobe Photoshop 2026 | Adobe | https://www.adobe.com/products/photoshop.html |
| **Other** |  |  |
| RC2 Rodent Circuit Controller | VetEquip | 922100 |
| Induction Chamber, 7 liter | VetEquip | 941488 |
| Color coded, nylon-reinforced, 1/4" ID, conductive gas supply hose | VetEquip | 931503 |
| 12mm Nosecone and 14mm Nosecone | VetEquip | 921612 and 921614 |
| Anesthesia silicone mask, reusable, size 00, premature baby | Drager | MP02900 |
| NORMOTHERM™ INFRARED HEATER | Britz & Company | N/A |
| SomnoSuite® Low-Flow Anesthesia System | Kent Scientific Corporation | SS-01 |
| PhysioSuite® | Kent Scientific Corporation | PS-02 |
| IntelliVue MX500 Patient Monitor | Philips Medizin Systeme Böblingen GmbH | 866064 |
| Micro NeoLead, AAMI radio lead | Philips Medical Systems Hsg | 989803183141 |
| Nasal Alar SpO2 Sensor | Philips Medical Systems Hsg | 989803205381 |
| NIBP Cuffs (Neonatal single-patient cuff size #1) | Philips Medical Systems Hsg | M1866B |
| Symmetry Pro-120 Multi-Pupose Electrosurgical Generator | Bovie | A1250S |
| Push Button Electrosurgical Pencil 50/Box | Bovie | ESP1 |
| Reusable Grounding Cable | Bovie | A1252C |
| Disposable Split Adult Return Electrodes | Bovie | ESRE-1 |
| 23 gauge, Hubless Needle, custom length (2 in), point style 4, 12 DEG, 6/PK | Hamilton Company | 22023-01 |
| 24 gauge, Hubless Needle, custom length (2 in), point style 4, 12 DEG, 6/PK | Hamilton Company | 22024-01 |
| 33 gauge, Small Hub RN Needle, custom length (2.5 in), point style 4, 12 DEG, 6/PK | Hamilton Company | 7803-05 |
| 5 µL, Model 75 RN Syringe, Needle Sold Separately | Hamilton Company | 7634-01 |
| 10 µL, Model 701 RN Syringe, Needle Sold Separately | Hamilton Company | 7635-01 |
| 50 µL, Model 710 RN Syringe, Needle Sold Separately | Hamilton Company | 7637-01 |
| 100 µL, Model 710 RN Syringe, Needle Sold Separately | Hamilton Company | 7638-017634-01 |
| Curved hemostatic forceps | Multiple sources | N/A |
| Ball end hex driver 5/64" | Multiple sources | N/A |
| Ball End Hex Driver 3/32" | Multiple sources | N/A |
| Pipette tips 20 µL, filtered | Multiple sources | N/A |
| Pipette tips 100 µL, filtered | Multiple sources | N/A |
| Pipette, single-channel, volume 20 µL | Multiple sources | N/A |
| Pipette, single-channel, volume 100 µL | Multiple sources | N/A |
| Palladium™ "Pebax™" Heat Shrink Tubing | Component Supply | PBST2-040-40-004C7635-01 |
| 1500 Watt 10 Amp 12 Temperature Heat Gun | Harborfreight | 69343 |
| 18.70 mm A/P Bar | KOPF | N/A |
| A.P. Slide Attachment | KOPF | Model 1261 A/P Slide Attachment |
| Rotation Adapter | KOPF | Model 1460-G Rotation Adapter |
| Travel Vertical Translation Stage | KOPF | VAP4/M |
| Stereotaxic arm and micro manipulator with fine adjustment | KOPF | Model 1460-61 |
| Single Axis Translation Stage | THORLABS | LT1/M |
| Cube Geared Head | Arca-Swiss | 8501303.1 |
| Aluminum Breadboard | THORLABS | MB2530/M |
| Animal cradle | NIMH/SI | N/A |
| Stereotaxic arm base holder | NIMH/SI | N/A |
| Ultrasound probe holder | NIMH/SI | N/A |
| Vevo MD Imaging System | Fujifilm Sonosite, Inc. | 51475 |
| UHF22 Transducer | Fujifilm Sonosite, Inc. | 51414 |
| UHF48 Transducer | Fujifilm Sonosite, Inc. | 51415 |
| Hair trimmer | WAHL | 41590-0438 |
| Hair trimmer blade | WAHL | 41590-7840 |
| Heat Pad - Large | Arbor Scientific | P3-1015-03 |
| 2x2 Gauze | Multiple sources | N/A |
| 4x4 Gauze | Multiple sources | N/A |
| Tongue depressors | Multiple sources | N/A |
| Sterile tongue depressors | Multiple sources | N/A |
| Cotton Swab | Multiple sources | N/A |
| Sterile cotton swab | Multiple sources | N/A |
| Tape | 3M | 1527-1 |

### Materials and equipment

Hormone stock solution:

| **Reagent** | **Final concentration** | **Amount** |
|---|---|---|
| [des-Gly10, D-Ala6]-LH-RH ethylamide acetate salt hydrate | 4 mg/mL | 5 mg |
| 1X PBS | -- | Up to 1.25 mL |
| Total |  | 1.25 mL |

Dissolve the hormone in 1x PBS, aliquot the stock solution, and store at -20°C. Hormone working solution:

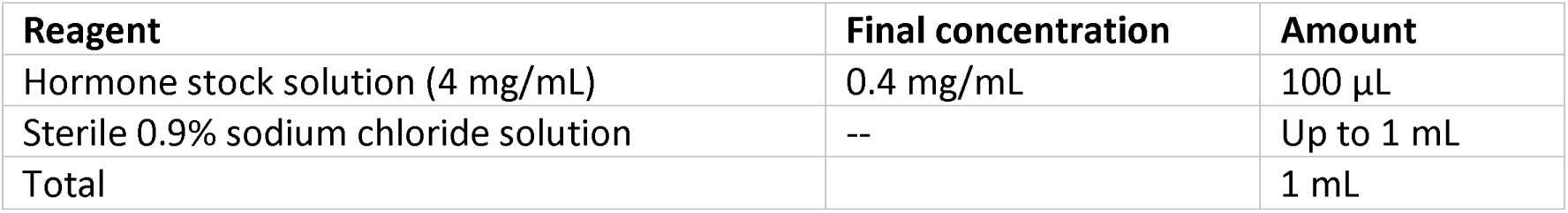

| **Reagent** | **Final concentration** | **Amount** |
|---|---|---|
| Hormone stock solution (4 mg/mL) | 0.4 mg/mL | 100 µL |
| Sterile 0.9% sodium chloride solution | -- | Up to 1 mL |
| Total |  | 1 mL |

Prepare fresh on the day of use. Thaw an aliquot at 4°C and dilute the stock solution with sterile 0.9% sodium chloride solution. Discard any unused working solution after use.

Key materials and equipment required for the protocol are shown in **Figures 1 and 2** and listed in the Key resources table. Drawings of the guide tube holder and cradle are available in **Data S1**.

**Figure 1.**
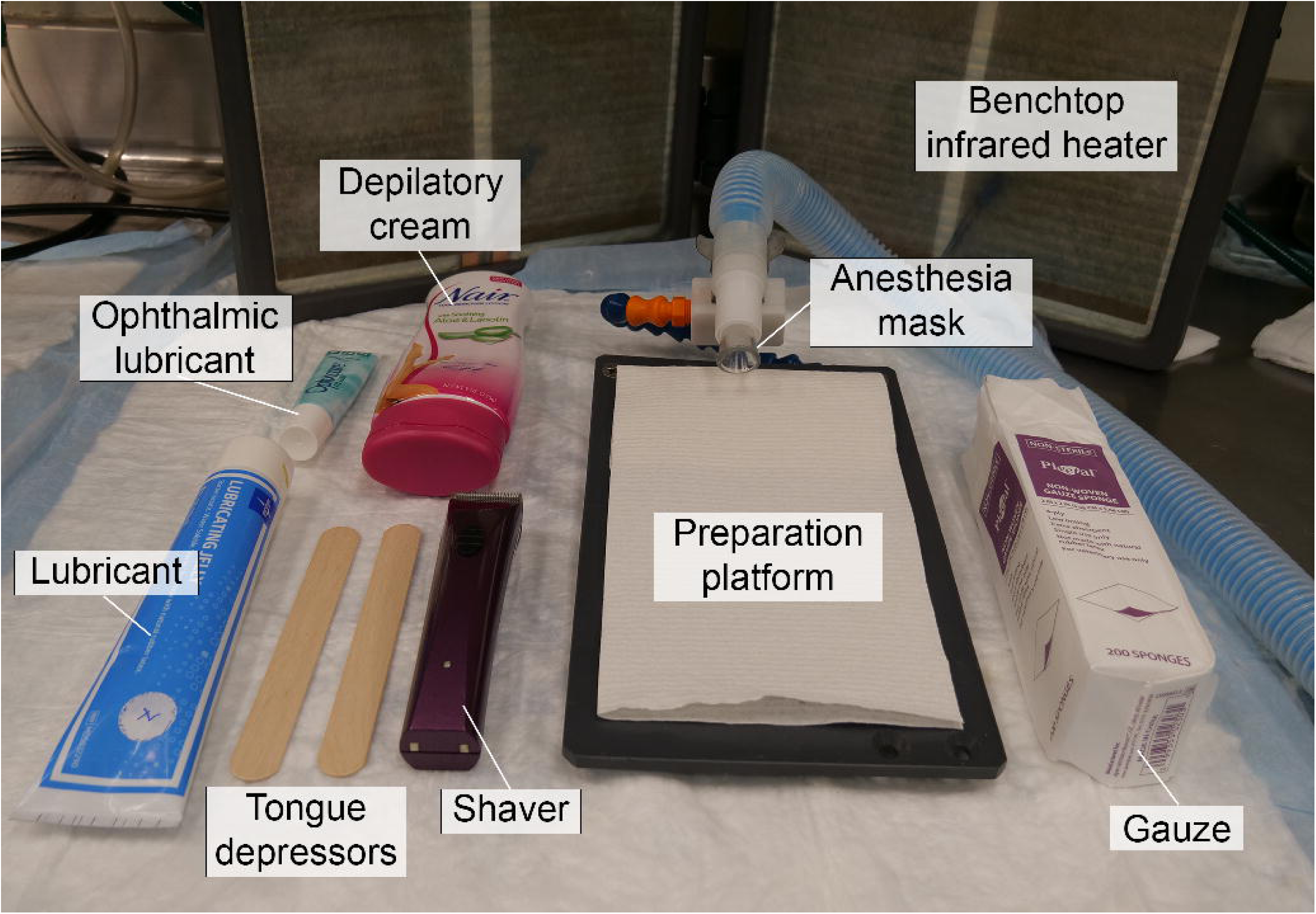
Materials and setup for preparation of the abdominal region of the dam. The preparation area includes a platform with support for stabilizing the anesthesia tubing over the dam’s mouth and/or nose while in the supine position. Tools and consumables for ophthalmic lubrication, trimming, and, when necessary, depilating the abdominal fur or hair are kept readily available. Following abdominal preparation, the animal is repositioned into the prone position for removal of a small dorsal fur patch to prepare for return electrode placement at the FIVI platform.

**Figure 2.**
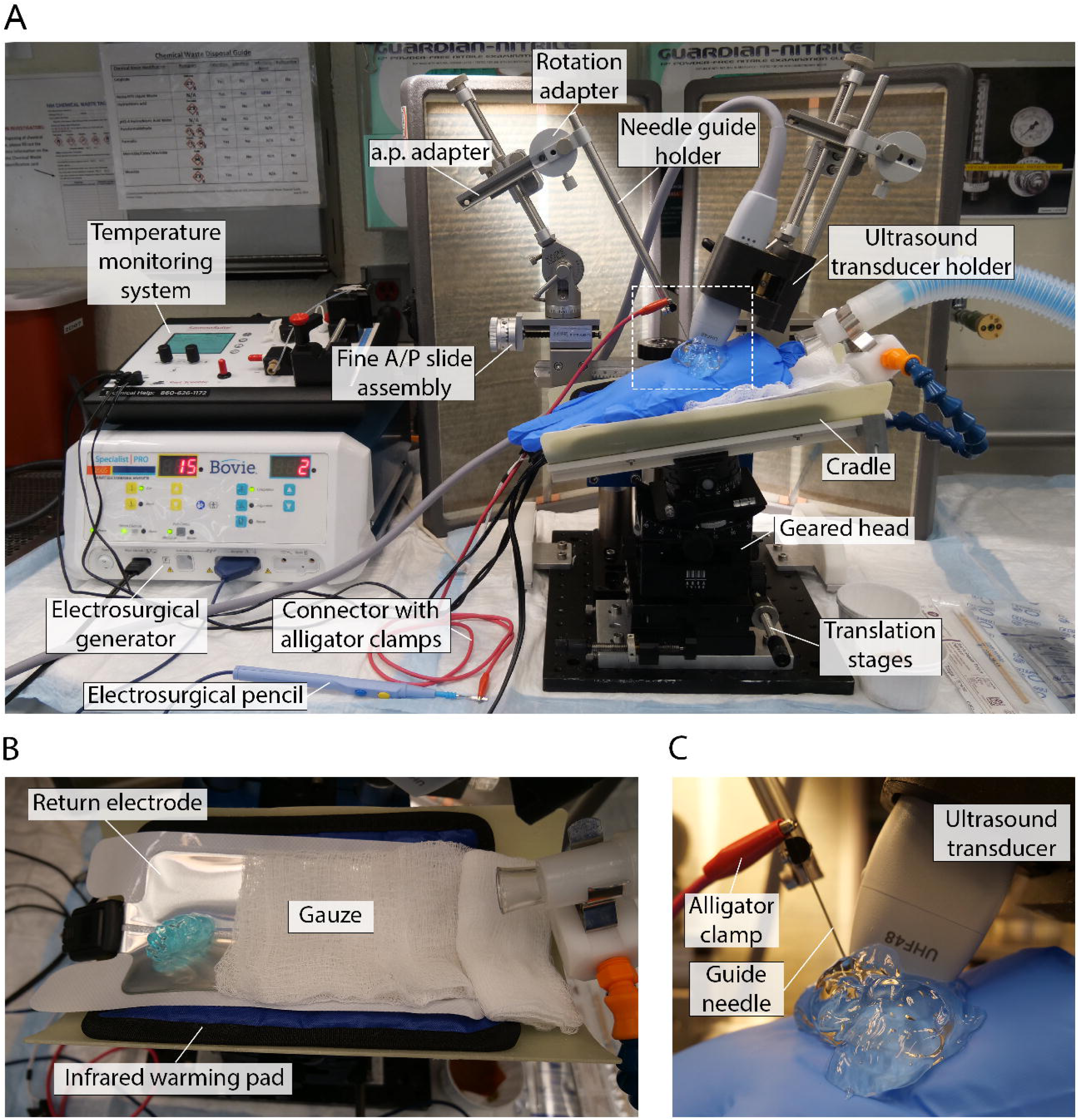
Materials and experimental setup for ultrasound-guided fetal intracerebroventricular injection (FIVI). The FIVI experimental area includes the injection platform, heating elements, physiological monitoring system, and electrosurgical unit (panel A). Following transfer from the preparation platform, the pregnant dam is positioned supine on the FIVI cradle over the return electrode (panel B). The electrode is covered with gauze except at the contact site with the shaved dorsal patch, and conductive gel is applied to ensure proper current return and minimize tissue injury. After stabilization in the cradle, the sterile field is prepared, and sterile ultrasound gel is applied over the abdomen. Only sterile or disinfected items contact the sterile field, including cotton-tipped applicators, tongue depressors, the guide tube and injection needle, and the ultrasound transducer. Using the translation stages and geared head, the target fetus is positioned beneath the ultrasound transducer. The fine anterior/posterior (A/P) slide assembly and A/P adapters allow precise positioning of the guide tube and transducer during fetal targeting. Once aligned with the transducer, the guide tube is advanced through the ultrasound gel toward the skin (panel C). Electrocautery-assisted entry is achieved through an alligator clamp connected to the exposed non-insulated portion of the guide tube (panel C), enabling skin penetration with minimal to no fetal displacement (see also Methods Video 2).

Once received, clean the custom-made guide needles (hubless Hamilton Company needles) with acetone followed by 70% ethanol. Once dried, sheath the guide needles with Palladium Pebax Heat Shrink Tubing. **Figure 3** shows the materials used.

**Figure 3:**
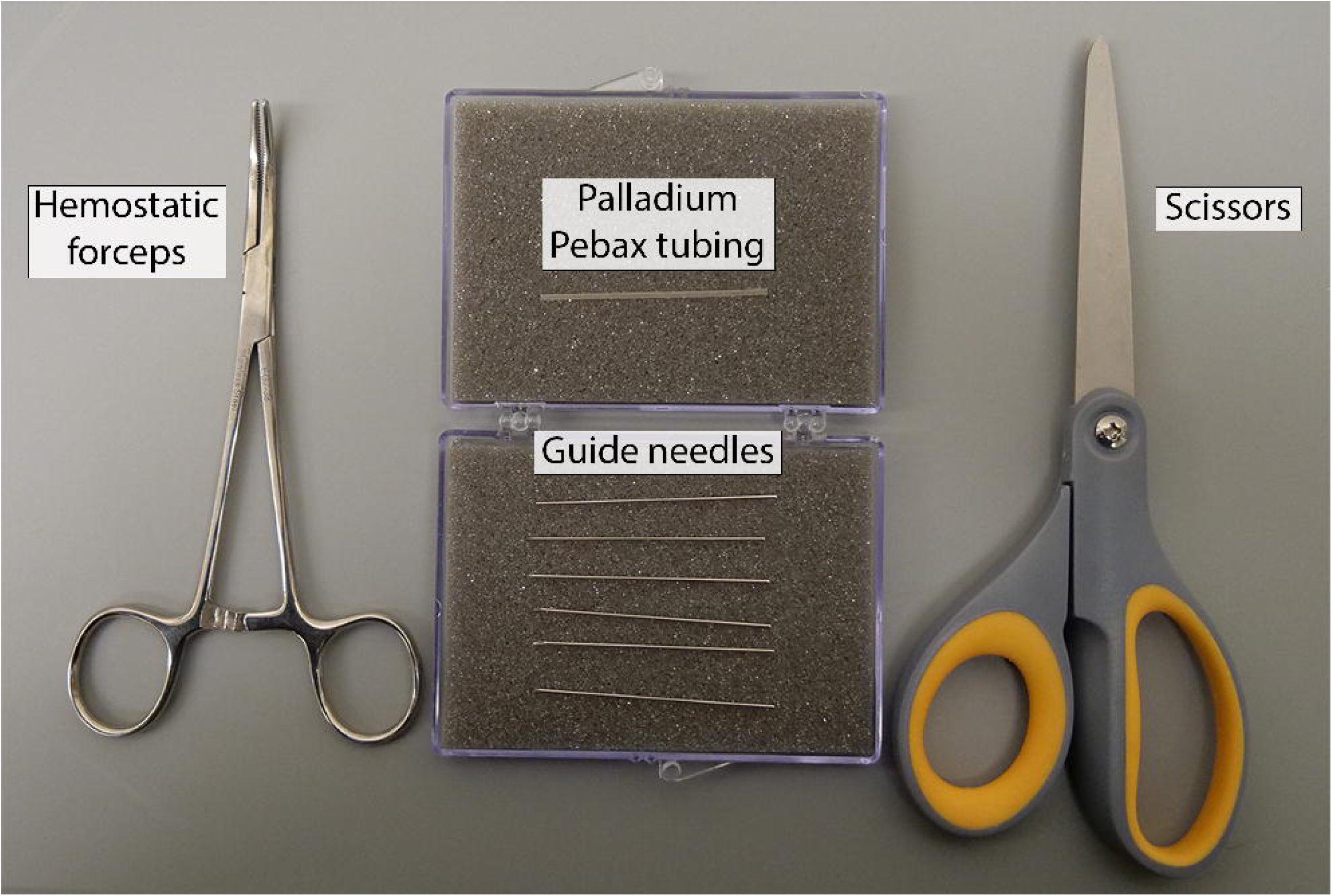
Materials used for sheathing guide tube needles.

Sheath guide needles using the following steps (see **Methods Video 1**):

1. Cut the Pebax tubing to a length of approximately 46 mm (∼1.8 in).
2. Position the tubing over the guide tube, leaving approximately 3 mm (∼0.12 in) of the needle exposed at the blunt end (**Figure 4A**). Maintain a distance of 1-2 cm between the heat source and the tubing during heating.
3. Using a heat gun (temperature setting: 300°C or 570°F), manually hold the guide needle and tubing, apply heat to the beveled end of the needle until the tubing is securely attached (**Figure 4B**). Ensure that the tubing is positioned immediately below the bevel to minimize current loss during cauterization.
4. Once enough unsheathed needle length is available at the blunt end to securely grasp the needle (**Figure 4C**):

i. Hold the needle with hemostatic forceps.
ii. Gradually move the heat source from the beveled end toward the blunt end over approximately 10-20 seconds
iii. Continue applying heat until the tubing is fully contracted around the needle with hemostatic forceps (**Methods Video 1**; **Figure 4D**). This allows the tubing to contract uniformly along the length of the needle and prevents the formation of uncoated segments that may impair current conduction through the needle.

**Figure 4:**
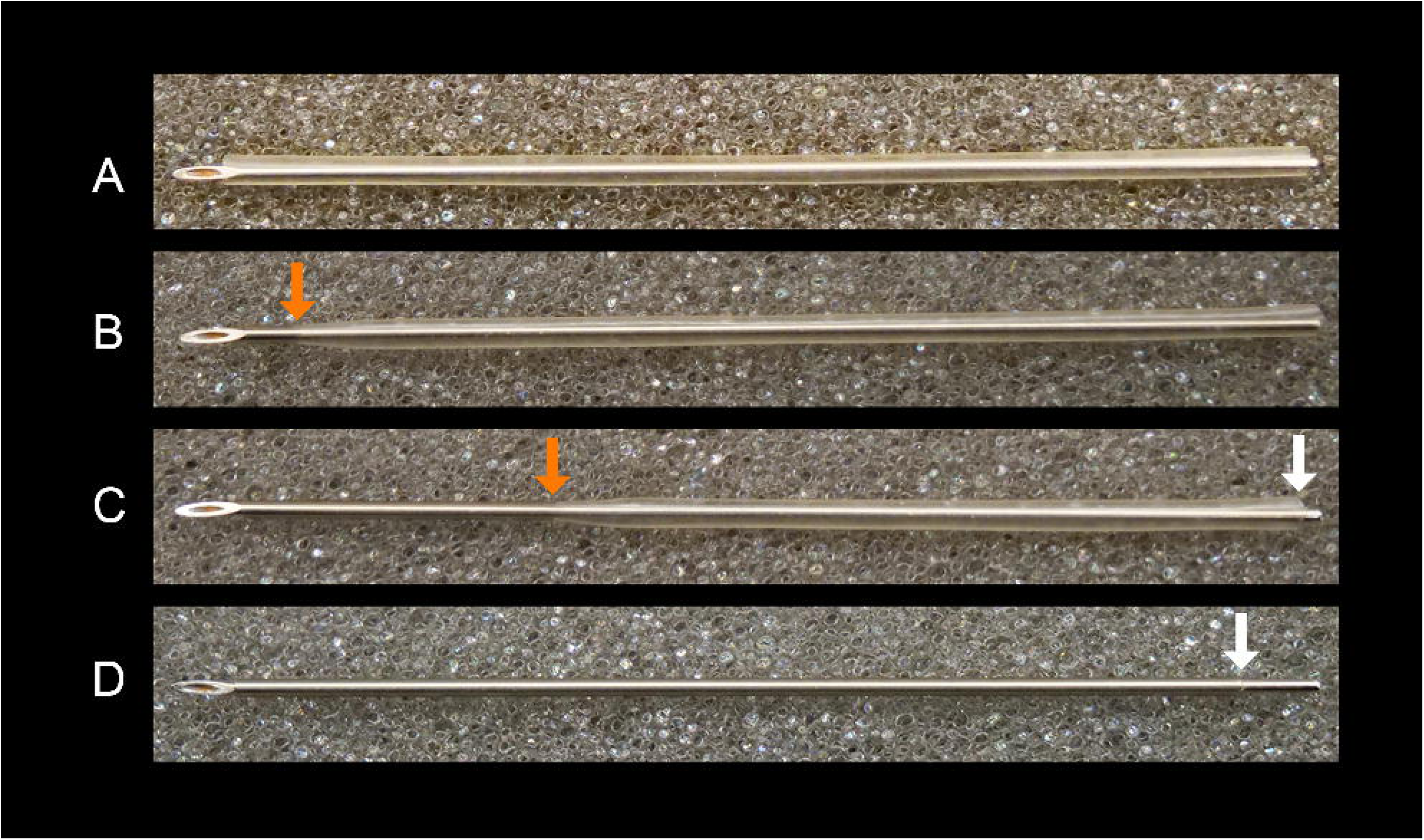
Palladium Pebax tubing contraction during guide needle sheathing. Panel A shows positioning of the tubing over the guide needle, leaving approximately 3 mm (∼0.12 in) of the needle exposed at the beveled end. Panel B shows initial contraction of the tubing beneath the bevel following heat application with a heat gun. Panel C shows further contraction of the tubing along the beveled end, resulting in retraction of the tubing from the blunt end toward the beveled end. Panel D shows the guide needle after complete tubing contraction. In panels B and C, orange arrows indicate the transition between contracted and non-contracted tubing. In panels C and D, white arrows indicate the tubing edge at the blunt end, showing progressive tubing retraction and the remaining exposed segment used for hemostatic forceps handling in C and alligator clamp attachment in D.

**Alternatives:** The Clarius L20 HD3 portable handheld ultrasound system can be used as an alternative to the Vevo MD ultrasound system (compare **Methods Videos 2** and **3**). Although image resolution is lower than that of the UHF48 transducer, which we use for most injections, and slightly lower than that of the UHF22 transducer, the device provides sufficient image quality for pregnancy confirmation, gestational staging, and FIVI. Its substantially lower cost may facilitate adoption of this protocol by a broader range of research groups.

**Alternatives:** To further reduce costs, the vertical translation stage (VAP4/M, KOPF) carrying the A/P bar (KOPF) can be omitted. Although this component provides additional flexibility for adjusting the distance between the dam and the guide or ultrasound transducer holders, these adjustments are infrequently required. Instead, the A/P bar can be mounted directly onto longer metal rods attached to the breadboard. If additional vertical distance is required, a longer metal rod can be used to carry the guide holder and ultrasound transducer.

## Step-by-step method details

### Preparation of pregnant dams for FIVI procedure

This section describes all pre-procedural preparation steps necessary for efficiently and safely perform the FIVI procedure, including: (1) transport to the procedure room and fasting (marmosets only); (2) anesthesia induction and maintenance; (3) physiological monitoring setup; (4) administration of analgesia and perioperative medications; (5) hair removal to improve echogenicity; (6) return electrode preparation; (7) positioning of the dam; and (8) sterile field creation.

#### Timing: 20-30 min

1. Set up the room with the tools, consumables, preparation and injection setups, and ultrasound machine (**Figures 1 and 2**).

2. Turn on the support heating elements before transferring the pregnant dam to the procedure room.

***Note:*** For both marmosets and rats, we used multiple heat sources while preparing the pregnant dam, including an infrared warming pad, a benchtop infrared heater and an overhead heating lamp. Continuous heating was provided by the warming pad, with supplemental heat from the other sources applied as needed to maintain body temperature between 37-38 °C. The infrared warming pad in the cradle is set between 40-42°C to compensate for heat loss between the pad and the animal. The benchtop infrared heater and overhead heating lamp do not have adjustable temperature settings and are therefore positioned according to the dam’s body temperature rather than at a fixed distance. Body temperature is continuously monitored using a temperature probe (see Step 15), and the position of the supplemental heat sources is adjusted as needed to maintain the target temperature range.

3. Transfer the pregnant dam from the housing room to the procedure room.

**CRITICAL:** For marmosets, perform the procedure after a fasting period of at least 4 hours to minimize the risk of regurgitation and aspiration during anesthesia. This timing might change in accordance with the institutional guidelines and facility-specific standard operating procedures. In our case, most procedures were performed following overnight fasting (approximately between 12-14 hours) without complications.

4. Weigh the pregnant dam and administer premedication as appropriate.

i. For marmosets, administer diazepam (1.0 mg/kg, 5 mg/mL; intramuscular) as a sedative and muscle relaxant, and atropine (0.2 mg/kg, intramuscular) to reduce salivary/respiratory secretions. Allow 10-15 min for absorption prior to induction of anesthesia. During this period, monitor for marked reduction in responsiveness, changes in breathing pattern or rate, and excessive salivation, which may be accompanied by coughing or whistling respiratory sounds.

***Note***: In this protocol, premedication is administered only to marmosets, but not to rats, consistent with the veterinary anesthetic protocols used for each species. Premedication requirements should follow institutional veterinary guidance, with doses adjusted based on the present of underlying clinical conditions and animal’s overall health.

***Note:*** Atropine is administered before anesthesia to reduce salivation. Although fetal cardiac complications have not been observed in our procedures, maternal atropine administration may affect fetal heart rate. Fetal cardiac activity should therefore be monitored under ultrasound during the procedure, with particular attention to sudden or prominent changes in rate.

***Alternatives***: Alternative sedative or pre-anesthetic regimens approved by the relevant Institutional Animal Care and Use Committee (IACUC) may be used.

5. Induce anesthesia with 3% isoflurane delivered at a flow rate of 1 L/min.

***Note:*** For marmosets, induce anesthesia using a face mask (VetEquip Mask #3 or Mask #4). For rats, perform induction in an anesthesia induction chamber.

6. Once the dam is anesthetized, transfer the animal to the preparation setup (**Figure 1**) and place it in the supine position. Monitor respiratory rate (rat: 70-90 breaths/minutes, marmoset: 30-50 breaths/minutes) and assess the absence of toe pinch reflex to confirm adequate depth of anesthesia.

***Alternatives***: A reusable heat pack (e.g., Arbor Scientific, see Key resources table) can be used as a supplemental heat source to help maintain the dam’s body temperature, in addition to a ceiling heating lamp or panel. Carefully monitor body temperature to prevent overheating and thermal injury to the dam.

7. Attach the anesthesia mask tubing to the anesthesia tubing holder in the preparation setup and deliver the anesthetic agent using species-specific mask configurations.

i. In marmosets, the mask (Drager anesthesia mask) is stabilized over the marmoset face, covering both mouth and nose.
ii. In rats, isoflurane is delivered through a fitted nose cone stabilized over the rat nose.

***Alternatives***: The procedure cradle may be used for both animal preparation and ultrasound-guided injection to maintain stable anesthesia and consistent temperature regulation.

8. Reduce the isoflurane concentration and maintain anesthesia at 1-2% for marmosets and and rats.

***Note:*** For both marmosets and rats begin maintenance anesthesia at approximately 2% isoflurane and titrate downward to the lowest effective concentration. An adequate plane of anesthesia is maintained when the animal shows no response to toe pinch while respiratory rate remains stable at an appropriate level. Avoid unnecessarily high isoflurane concentrations to minimize anesthetic exposure of the dam and fetuses. Maintaining the lowest effective isoflurane concentration generally facilitates faster recovery of the pregnant dam.

9. Set up the physiological monitoring system and initiate continuous monitoring of vital (cardiovascular, respiratory, and central nervous system) functions.

***Note:*** Use a physiological monitoring system to monitor pulse oximetry and regulate body temperature (marmoset: IntelliVue MX500; rat: Kent Scientific SomnoSuite or PhysioSuite). In marmosets, additionally monitor end-tidal CO_2_, respiratory rate, and heart rate. Provide supplemental heating using a benchtop infrared heater and an overhead heating lamp to maintain body temperature between 37-38°C to prevent hypothermia.

### CRITICAL

Monitor vital functions, and toe pinch reflex throughout the procedure and record values every 10 min. Manually assess respiratory rate to ensure appropriate isoflurane delivery and prevent anesthetic overdose. Assess proper mucous membrane color. At minimum, monitor temperature and SpO_2_ in rats and marmosets. In marmosets, we recommend additional monitoring of end-tidal CO_2_.

Adjust the isoflurane concentration as needed (e.g., in response to slow or rapid breathing), and pause the procedure until physiological parameters stabilize if necessary.

10. Administer perioperative medications.

i. In marmosets, administer meloxicam (0.1–0.2 mg/kg, 5 mg/mL; intramuscular or subcutaneous) for preemptive analgesia. Additionally, administer 3 mL of sterile 0.9% sodium chloride solution warmed to 37°C to help prevent dehydration.
ii. In rats, administer meloxicam (1 mg/kg; 1.5 mg/mL; intramuscular or subcutaneous) for preemptive analgesia.

***Alternatives:*** Other agents approved by the relevant IACUC may be used.

11. Apply ophthalmic ointment to both eyes to prevent corneal drying.

12. Trim the abdominal fur and apply depilatory cream to eliminate the undercoat. Apply the depilatory cream to the abdominal surface, immediately caudal to the last ribs. For later gestational stages, extend the application to the lateral aspects of the abdomen to facilitate access to fetuses positioned laterally.

### CRITICAL

When trimming the abdomen of the rat dam, pay special attention to the nipples to prevent injury. If needed, mark the placement of the needles with a pen for better retrieval. In both rat and marmosets, carefully control the application time of the depilatory cream to prevent chemical burns. Apply a thick layer of cream to the trimed area. After 30 seconds, begin removing the cream using a tongue depressor to assess hair removal. If necessary, reapply the cream in 10 segments until hair is fully removed. Remove the bulk of the cream with a tongue depressor, then gently clean the area with warm water or saline using a compress to eliminate all residues and prevent skin lesions. Do not use alcohol to remove the cream, as this may cause skin irritation.

***Note:*** In marmosets, depilatory cream is not always required. When needed, it is typically used only for the first procedure. In subsequent procedures, adequate echogenicity is usually achieved by trimming the fur alone.

13. Reposition the animal in the prone position and trim a small fur patch (approximately 2 x 3 cm) on the dorsal surface, anterior to the tail base, to allow placement of the cautery return electrode.

14. Remove all loose hair clippings and transfer the dam from the preparation area to the procedure cradle (**Figure 2A**). Position the animal in the supine position over the electrosurgical grounding electrode (**Figure 2B**). Because the electrode surface is similar in size to the animal, cover it with gauze, leaving exposed only the area that will contact the shaved dorsal skin to ensure effective grounding while minimizing unnecessary skin contact.

#### Note

Apply non-sterile ultrasound gel or lubricant between the trimmed dorsal area and the dispersive electrode to improve contact and ensure safe return of electrosurgical energy delivered through the guide.

15. Once the dam is stabilized on the electrosurgical grounding electrode, insert the rectal temperature probe and begin continuous temperature monitoring. Adjust supplemental heating as needed to maintain body temperature at 37-38°C.

#### Note

The SomnoSuite or PhysioSuite system is used for continuous temperature monitoring and control of the heating pad in both marmosets and rats.

16. Angle the cradle at 25-35° (geared head scale, where the vertical position is 90°) to elevate the head and reduce pressure on the inferior vena cava caused by the weight of the gravid uterus.

17. Create a sterile field by performing three alternating scrubs of the abdominal skin using 7.5% povidone-iodine solution (e.g., Betadine Surgical Scrub) and 70% ethanol. At each scrub, use a circular motion moving from the center outward. Proceed sequentially between alternating applications. After the final application, allow the skin to dry completely before proceeding.

### CRITICAL

When creating the sterile field, use only the minimum volume of antiseptic solution necessary to avoid soaking the fur. Excess moisture can contribute to heat loss and hypothermia. Before initiating ultrasound-guided injections, confirm that physiological parameters have stabilized. After establishing the sterile field, ensure that only sterile or disinfected instruments come into contact with the prepared skin. These include the sterile guide, disinfected ultrasound transducer, and sterile tongue depressors or cotton-tipped applicators used to apply sterile gel or remove air bubbles as needed. The operator may manipulate components outside the sterile field, such as the stereotaxic arm and support rods, using clean gloves, but should not directly contact the prepared skin or sterile components that contact the animal.

### Ultrasound-guided fetal injections

This section outlines the ultrasound-guided fetal injection procedure. The protocol combines stereotaxic mechanical control with real-time ultrasound imaging to enable selective targeting of the fetal brain. It describes the steps involved in (1) injection planning, (2) fetal targeting, and (3) controlled tissue penetration, along with troubleshooting guidance. Injection trajectory and depth are determined individually for each fetus under real-time ultrasound visualization, regardless of species, based on gestational stage, fetal position, and target accessibility. When possible, plan for two operators to perform the procedure. While one operator identifies the fetuses and plans the trajectory, the second operator can monitor vials and prepare and load the syringe. This approach reduces the time required per injection and shortens the overall procedure duration.

#### Timing: 5-20 min (per fetus)

18. Disinfect the ultrasound transducer by draping the transducer in sodium hypochlorite-based disinfectant wipes (e.g. Sani-Cloth Bleach, orange top) and allow it to remain in contact for 4 min to ensure sterilization.

#### Note

Sodium hypochlorite-based disinfectant wipes were used with the UHF22 and UHF48 transducers in this protocol. Transducer manufacturers may recommend multiple compatible disinfectants. Therefore, users should follow the manufacturer’s guidelines to ensure compatibility of the disinfectant with the transducer.

19. Secure the ultrasound transducer in its custom-designed holder (**Figure 2A**).

20. Apply sterile ultrasound gel to the abdomen. Use a sterile tongue depressor to spread the gel evenly as needed, ensuring a layer thickness of approximately 1-2 cm to allow proper immersion of the angled transducer (**Figure 2C**).

#### Note

To minimize heat loss, the ultrasound gel can be warmed to 37°C (e.g., in incubator or warming pad) prior to application. Gel warming is particularly useful when additional heat loss could compromise maintenance of the dam’s body temperature. Continuously monitor body temperature and adjust the available heating sources as needed to maintain the target range of 37-38°C.

21. Position the angled ultrasound transducer against the abdominal skin within the sterile ultrasound gel layer (**Figure 2C**).

#### Note

the ultrasound transducer holder is carried by a rotator mounted on a stereotaxic arm. Position the ultrasound transducer at approximately 30-50° relative to the abdominal skin surface to provide sufficient clearance for the guide needle (**Figure 2**). This range provides an initial working angle and can be adjusted during targeting using the rotator or stereotaxic arm to optimize visualization and alignment of the injection trajectory.

22. Mount the guide needle in the guide holder using sterile hemostatic forceps. Use the ball end hex driver 5/64" to loosen the screw holding the two guide holder plates and insert the guide needle between them.

#### Note

The angle of the arm carrying the guide holder depends on the planned trajectory but is typically set between 35-45° to allow for the guide needle beneath the transducer (**Figure 2C**). The rotation adapter should be set at 0°.

23. Use the ultrasound transducer and the mechanical translation plates supporting the cradle to position the target fetus for injection. Use the geared head to adjust the dam’s position as needed to optimize fetal alignment for targeting. If visualization of the fetus is poor, see Troubleshooting, Problem 1.

24. Advance the guide needle through the ultrasound gel and align it to the ultrasound transducer over the fetal head (**Figure 2C**).

25. Adjust the angle of the guide tube by modifying the stereotaxic arm to align with the planned injection trajectory and optimize fetal targeting.

### CRITICAL

When determining the optimal injection trajectory, avoid nipples (in rats), maternal internal organs (e.g., bladder and intestines), and large vessels (e.g., umbilical cord). Whenever possible, avoid penetrating the placenta by repositioning the pregnant dam using the geared head and translation plates. If placental penetration is unavoidable for proper targeting, use color Doppler mode on the ultrasound system to identify major vessels and minimize the risk of hemorrhage along the needle trajectory. For most injection penetrations done through the placenta, no blood is observed upon withdrawal of the needle and guide. In rare cases, a minimal amount of blood may be observed during needle retraction, without evidence of active bleeding. If the placenta or umbilical cord obstructs the injection trajectory, see Troubleshooting, Problem 2.

26. While gently pressing the guide needle against the skin, apply a brief electric current (1-3 seconds) to the non-insulated blunt side of the guide to help tissue penetration with minimal displacement of the fetus (**Methods Video 4**).

#### Note

Because the 12° bevel angle of the injection needle allows penetrating the abdominal wall and uterus, advancing the guide tube beyond the subcutaneous tissue layer is not necessary. We have tested injection needles with 12°, 30° and 60° bevel angles and observed smoother tissue penetration with the steeper angle. Use monopolar cut mode at 15 W. Deliver the current to the guide through a connector with an alligator clamp attached to the non-insulated blunt end of the guide and activate the current using the electrosurgical pencil (**Figure 2A**).

#### Note

If air bubbles generated during electrocautery reduce ultrasound visualization, see Troubleshooting, Problem 1.

27. Load rAAV vector preparation into the Hamilton syringe.

#### Note

High levels of viral transduction are achieved using high-titer preparations (e.g., Addgene rAAV viral preparations ≥ 1x10 vg/mL). The injected volume varies depending on gestational age at the time of injection and the construct (e.g., for rAAV maximum of 60 μL per fetus in marmosets and 10μL in rats lead to broad and dense transduction). Injection parameters should be empirically optimized for each experimental condition (e.g., viral vector, gestation stage) and species.

28. Smoothly advance the injection needle manually through the guide, penetrating the abdominal wall and uterine muscle, and continue into the uterine lumen adjacent to the fetal skull under ultrasound guidance.

#### Note

With experience, manual advancement of the injection needle provides tactile feedback when traversing the different abdominal and uterine tissue layers. This sensory feedback can also assist in assessing the relative firmness of the developing fetal skull prior to injection.

#### Note

If the needle angle or trajectory requires correction after tissue entry, see Troubleshooting, Problem 3.

29. Under ultrasound guidance, manually penetrate the fetal skull using a brief controlled jolt that advances the needle tip by approximately 1-3 mm (**Methods Video 5**, see timeline 1 to 2 seconds). Injection into the intended ventricular target is confirmed by visualizing the needle tip under ultrasound before vector delivery.

#### Note

If penetration of the skull is unsuccessful, see Troubleshooting, Problem 3.

30. Manually inject the viral vector into the target region under ultrasound guidance using repeated, small depressions of the syringe plunger. Deliver the full volume incrementally rather than as a single bolus, generally over approximately 30 seconds.

#### Note

When targeting ventricles located close to the fetal skull (e.g., the lateral ventricles), we recommend a longer dwell time before needle withdrawal, up to 5 minutes. The minimum dwell time required to prevent backflow has not been systematically evaluated, and backflow cannot be reliably assessed by ultrasound during the procedure. Confirm successful targeting by visualizing the needle tip position under ultrasound. Introducing a small air bubble at the end of the injection can increase the echogenicity of the injectate, thereby facilitating visualization and confirmation of the target site.

31. Withdraw the injection needle from the fetus and dam under ultrasound guidance.

32. Remove the guide tube from the dam skin.

33. Assess fetal viability by monitoring fetal cardiac activity under ultrasound to confirm stable heart function. Confirm the presence of a regular heartbeat and monitor for sudden or prominent changes in cardiac activity during and following the injection.

#### Note

Fetal heart rate is not routinely quantified during the procedure. Cardiac activity is assessed qualitatively under ultrasound, with particular attention to sudden or prominent changes relative to the activity observed in the individual fetus before injection. If sudden or prominent changes in fetal cardiac activity are observed following injections, see Troubleshooting, Problem 4.

34. Repeat steps 23 to 33 with the other fetuses.

#### Note

Unless bent, the same guide needle can be reused across injections in different fetuses. It may also be reused across procedures following thorough cleaning with acetone and subsequent disinfection with 70% ethanol. The procedure results in minimal to no displacement of the fetuses. Injected fetuses can therefore be distinguished from non-injected fetuses based on their position and orientation within the abdomen. The maternal skin entry sites associated with the previous injection trajectories provide additional landmarks for identifying fetuses that have already been injected.

### Post-procedure monitoring and recovery

This section outlines the post-procedural care steps following completion of the ultrasound-guided injection procedure.

#### Timing: 15-25 min

35. Remove the ultrasound gel from the abdominal surface using sterile gauze.

36. Apply a thin layer of local anesthetic (e.g., lidocaine ointment USP, 5%) sufficient to cover each skin entry site to reduce discomfort. followed by a thin layer of triple antibiotic ointment over the same area to minimize the risk of infection. Avoid excess application to minimize the risk of lidocaine toxicity.

37. Allow the pregnant dam to recover in a warm environment prior to returning it to the housing room.

38. Once the animal is fully awake, upright, and responsive to its surroundings, administer a small volume of 50% dextrose (marmoset: 0.1-0.2 mL, oral transmucosal; rat: 0.1 mL, oral transmucosal) as needed to prevent or treat hypoglycemia and support post-procedural recovery. Transfer the dam to the housing room only after full recovery is confirmed.

#### Note

In marmosets, monitor animals at least once daily for 7 days following the intrauterine injection procedure for signs of abdominal pain, uterine contractions, or vaginal discharge.

### Postnatal identification of transduced animals

This section outlines the procedures used for in vivo identification of transduced animals from birth.

#### Timing: 10-20 min

39. In a dark environment, use a fluorescence flashlight equipped with high-intensity LEDs and the appropriate excitation filter set to detect reporter expression through the skull and/or skin. For green-shifted reporters, an option is the model Xite-RB (Nightsea) with a Royal Blue excitation filter (440–460 nm) and 500–560 nm bandpass barrier filter glasses. For red-shifted reporters, model Xite-GR (Nightsea) with a Green excitation filter (510–540 nm) and 600 nm bandpass barrier filter glasses.

#### Note

Transgene expression persists at high levels into adulthood and remains detectable in hairless regions. However, identification of transduced animals is most reliable during the perinatal period.

#### Note

If there is reduced fetal survival following injection, see Troubleshooting, Problem 5. If reporter expression is not readily detectable at birth, see Troubleshooting, Problem 6. If post-procedural analysis reveals uneven or asymmetric transduction, see Troubleshooting, Problem 7.

## Expected outcomes

FIVI can be successfully performed across a broad gestational window. In marmosets we performed FIVI from PC62 to PC121 and in rats from PC13 to PC21. Procedural considerations vary with gestational age. At earlier stages, the smaller fetus has greater mobility within the uterine lumen, which may make stabilization and targeting more challenging, whereas at later stages, fetal size and position may restrict access to some fetuses or injection trajectories. Successful procedures are characterized by stable maternal and fetal physiological parameters throughout the intervention, maintenance of fetal cardiac activity following injection, and rapid recovery of the dam. In our experience, pregnancies proceed to term within the expected gestational timeframe, and offspring are delivered without complications, such as dystocia. Only rarely has FIVI resulted in miscarriage. Further discussion of procedural outcomes is provided in Ribeiro Gomes et al. (2026).

**Figure 5** shows successful transduction following FIVI performed at two developmental stages, PC83 and PC94. The representative examples shown here were generated using rAAV2, but similar widespread transduction patterns have also been observed using AAV9 and AAV8 (Ribeiro Gomes et al., 2026). This highlights a key feature of the approach: broad transduction is a general result of fetal CSF delivery rather than a property unique to a specific rAAV serotype.

**Figure 5.**
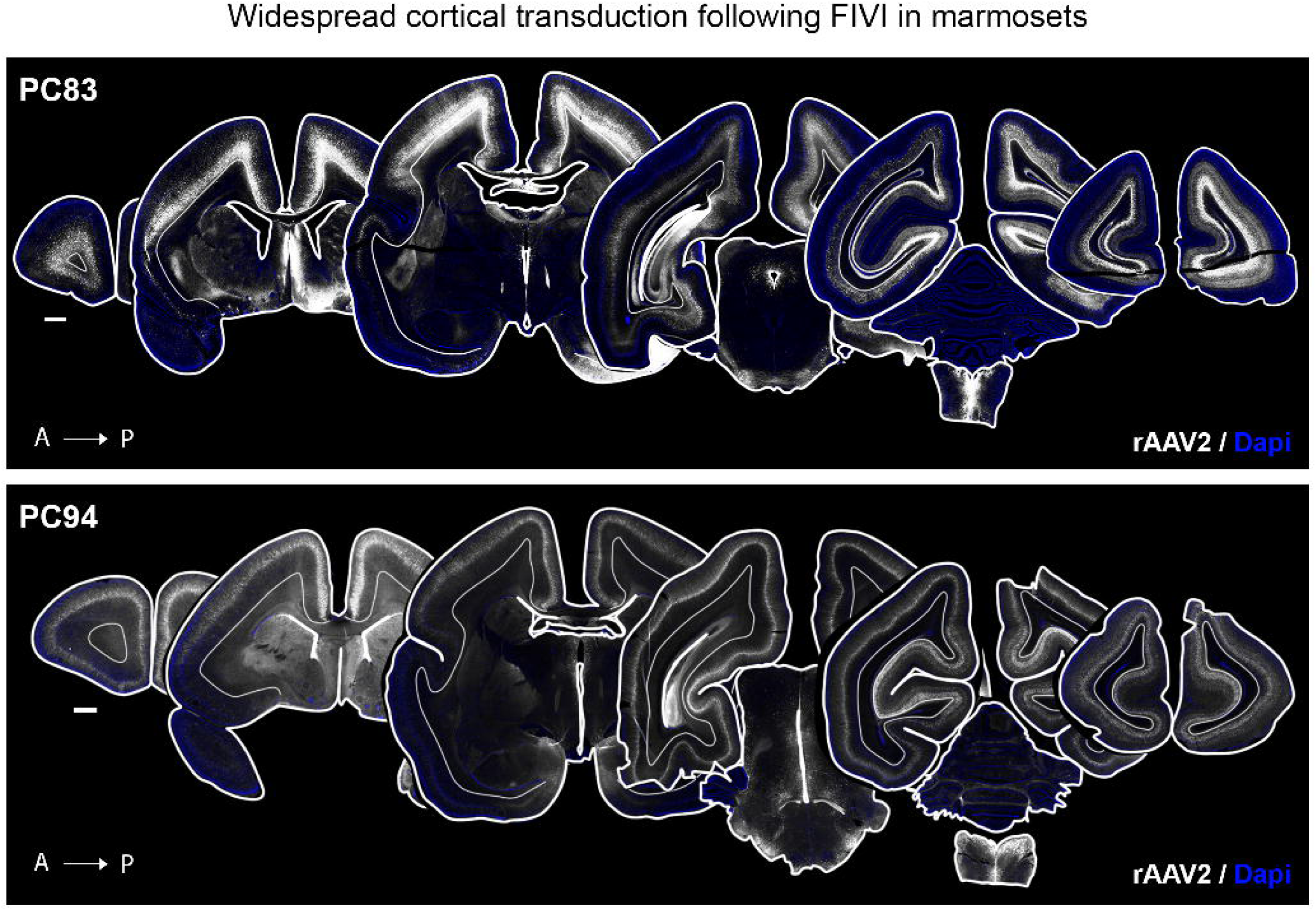
Fetal intracerebral delivery of rAAV leads to extensive transduction of cortical neurons in marmosets. Representative marmoset coronal sections illustrating extensive cortical transduction across anterior-to-posterior regions following FIVI performed at two developmental stages and analyzed at postnatal day 0. Broad cortical transduction is achieved following injections at both gestational ages, demonstrating the feasibility of FIVI-mediated gene delivery across developmental windows. The examples shown are from rAAV2 FIVI. FIVI volume/titer: 40 μL/8.6×10^12^ at PC83; 40 μL/1.5×10^13^ at PC94.. Scale bars: 1 mm.

Another key feature of rAAV delivery into fetal marmosets is the increased temporal resolution for cell targeting afforded by their prolonged gestation (143 days). **Figure 6** illustrates how injection timing influences transduction patterns, with earlier injections preferentially labeling neurons in deeper cortical layers and later injections labeling progressively more superficial populations. Cell populations permissive to rAAV transduction at earlier developmental stages are no longer efficiently transduced at later time points. Although the mechanisms underlying this developmental tropism remain unclear, our working hypothesis is that transduction efficiency depends on the maturation state of the targeted cells. Similar gestation-dependent patterns are observed in rats, although they are less pronounced because of the substantially shorter gestational period (23 days).

**Figure 6.**
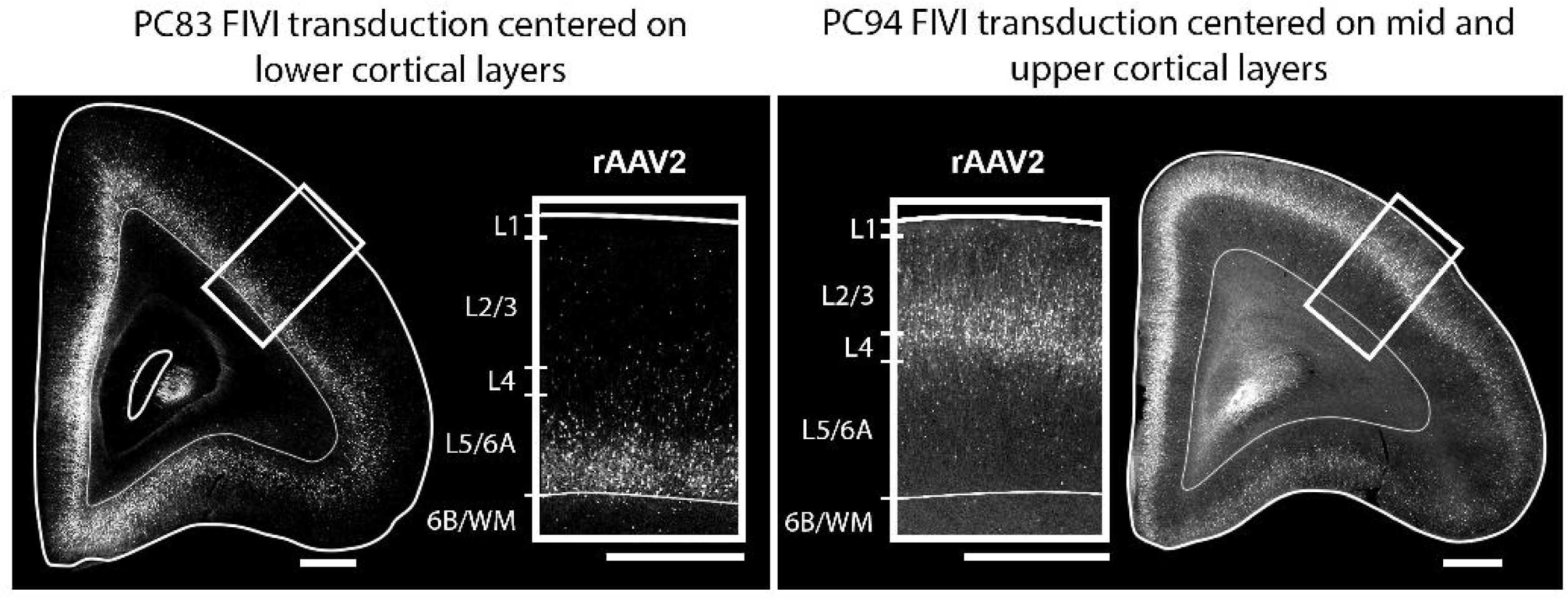
Gestational age influences transduction patterns following FIVI-mediated delivery of rAAV. Higher-magnification views of the prefrontal cortex from the animals shown in Figure 5. Transduced cells following PC83 FIVI are predominantly located in layers 5 and 6, whereas cells transduced following PC94 FIVI are predominantly located in layers 2 to 4. These differences illustrate how the prolonged gestation of marmosets enables temporal control of transgene delivery and cell-type targeting.

Further discussion of these developmental transduction patterns and additional strategies for targeting cell populations are provided in Ribeiro Gomes et al. (2026).

### Limitations

#### Limitation 1

The number of embryos that can be successfully injected is constrained by intrauterine space and fetal positioning. In rat dams, the number of fetuses injected typically ranges from 6 to 9 per litter, depending on gestational stage and uterine accessibility. At earlier gestational ages, the uterine horns are frequently positioned beneath maternal internal organs, limiting visualization and access. At later gestational stages, reduced intra-abdominal space may result in superimposed fetuses or positioning beneath maternal organs, further restricting access. Additionally, ultrasound image quality decreases for deeper fetuses, which may prevent reliable targeting. Although rare, similar limitations are observed in marmosets during advanced pregnancies, where deeper fetal positioning can prevent injection of all fetuses within a litter. In both species, fetal orientation may also limit targeting. Although repositioning the pregnant dam using the mechanical degrees of freedom of the cradle often improves access, certain anatomical constraints, such as the presence of the umbilical cord within the planned injection trajectory, may require exclusion of specific fetuses to avoid injury. In rats, where litters are large, we have successfully injected up to 9 fetuses during a single procedure. In marmosets, we often inject the entire litter and have successfully injected litters of up to 4 fetuses. In both species, the number of fetuses successfully injected generally increases as the operator gains experience with fetal identification, trajectory planning, and needle positioning.

#### Limitation 2

The number of fetuses injected per procedure is also constrained by the procedure length. In both rat and marmoset dams, the total number of fetuses that can be injected is limited by the duration of anesthesia. To minimize potential adverse effects associated with prolonged anesthesia, including impacts on fetal development and maternal recovery, the total procedure time is restricted to 2 hours or less.

#### Limitation 3

Variability in viral transduction. Although efficient and widespread rAAV transduction can be achieved following FIVI, variability in distribution may occur. For instance, control of the vector backflow and dispersion outside of fetal skull is limited. In some cases, transduction may be weaker or more prominent unilaterally or relatively higher in the spinal cord than in the brain, depending on viral diffusion and injection dynamics.

#### Limitation 4

Direct quantitative comparisons of fetal survival, pregnancy outcomes, and maternal recovery between FIVI and open surgical fetal interventions in nonhuman primates are currently lacking. Thus, although FIVI avoids laparotomy, uterotomy, and direct manipulation of the uterus or fetuses, whether these procedural differences translate into improved maternal or fetal outcomes remains to be quantitatively established.

#### Limitation 5

This cross-species protocol was successfully applied to fetal injections in rats and marmosets. The step-by-step protocol could be adapted to additional species and targets but may require adjustments to specific components, such as injection needle length and gauge for deeper fetuses or harder tissues (e.g., more developed fetal skulls), and the dimensions, mounting configuration, and range of motion of the injection platform for smaller or larger animals.

## Troubleshooting

### Problem 1

Poor ultrasound visualization of the fetus (refer to steps 23 and 26).

### Potential solution

• Ensure adequate removal of the dam’s hair (e.g., trimming and/or depilation) to improve ultrasound image quality.

• Ensure sufficient sterile gel thickness (approximately 1-2 cm for angled transducer immersion) to cover the base of the transducer.

• Air bubbles may be introduced into the ultrasound gel during application or electrocautery-assisted skin penetration (**Methods Video 4**). Due to the angled transducer approach, these bubbles generally move outside the imaging field and cause minimal disruption. If visualization is impaired, keep the guide tube in place and retract the transducer using the z-axis of the stereotaxic arm. Use a sterile cotton-tipped applicator or tongue depressor to displace the bubbles from the imaging field, and then return the transducer to the original imaging position.

• Reposition the dam using the translation plates and geared head to improve fetal alignment.

• Reapply gel if necessary to restore acoustic coupling.

### Problem 2

Placenta or umbilical cord obstructing injection trajectory (refer to steps 25)

### Potential solution

• Use color Doppler mode to identify and avoid major vessels along the planned path.

• Reposition the dam using the geared head to displace the placenta or umbilical cord out of the field of view.

• Modify the trajectory by introducing the needle in the amniotic sac lumen near the injection target. Then use the translation stages or stereotaxic arm movement to slowly approach the needle to the target while displacing the vessel away from the field of view and planned trajectory. Do so without contacting the vessel with the injection needle bevel.

• When safe trajectory cannot be achieved without compromising a blood vessel, omit injection of that fetus to prevent injury. If transplacental injection is required, proceed with caution and closely monitor for bleeding under ultrasound visualization.

### Problem 3

Difficulty penetrating the fetal skull (refer to steps 28 and 29).

### Potential solution

• Approach the fetal skull at an angle that provides sufficient mechanical advantage for penetration and minimizes displacement of the fetal head away from the needle or outside the ultrasound field of view (**Methods Video 6**). If necessary, adjust the injection angle using the rotator supporting the guide tube holder.

• When possible, plan the injection trajectory through regions of thinner developing bone to facilitate penetration.

• If the trajectory is misaligned or skull penetration is unsuccessful, first attempt to realign the trajectory without removing the needle from the uterine lumen. Reposition the dam using the translation stages and geared head or adjust the injection needle angle using the rotator supporting the guide tube holder, while keeping the guide in place.

• If adequate realignment cannot be achieved with the needle within the uterine lumen, retract the needle to the subcutaneous space and readjust the trajectory before attempting penetration again. Complete removal of both the injection needle and guide tube is rarely required.

### Problem 4

Transient changes in fetal cardiac activity following injection (refer to step 33).

### Potential solution

• Transient, visually apparent changes in fetal cardiac activity were occasionally observed in rat fetuses at the earliest gestational stage tested (PC13). In these cases, cardiac activity spontaneously returned to the pre-injection pattern without intervention. Allow time for recovery before proceeding with additional injections.

• Reduce the injected volume if bradycardia or irregular cardiac activity is observed. In subsequent PC13 rat procedures, the injection volume was reduced from 10 µL to 5 µL, after which similar changes were no longer observed. Although this was not systematically evaluated, reducing the injection volume may help minimize transient changes in fetal cardiac activity at early gestational stages.

### Problem 5

Reduced fetal survival following injection (refer to step 39).

### Potential solution

- Optimize injected volume and viral titer according to gestational stage and viral construct. Excessive volume or high titer may increase physiological stress and reduce survival.
- Minimize total procedure time and ensure stable physiological parameters throughout the procedure (e.g., adjust isoflurane concentration to the lowest effective level for each animal to maintain stable anesthesia).
- Confirm proper trajectory planning to avoid major vessels and excessive tissue disruption.

### Problem 6

Difficulty identifying injected neonates (refer to step 39).

### Potential solution

• Reporter expression may not be readily detectable at birth depending on the viral construct (e.g., promoter strength), titer, injected volume, and anatomical distribution. In rats, detectable reporter expression is often most prominent in the brain, spinal cord, back musculature, and facial regions due to peripheral nerve labeling. In marmosets, expression is typically most readily observed in less pigmented and hairless regions.

• When reporter expression is expected to be difficult to detect at birth, consider co-injecting a complementary vector or serotype expressing a readily detectable reporter to facilitate postnatal identification in rats and marmosets. Co-injection with a vital dye, such as Fluorescein, Indigo Carmine, Evans blue, can also be used in rats.

• Intramuscular rAAV injection at the time of the fetal procedure can serve as an auxiliary marker to assist in retrieving injected neonates in both species. For intramuscular delivery, select a fetal muscle that can be safely accessed under ultrasound based on fetal position. We have tested injections into the marmoset arm muscle because of its accessibility. A 5 µL injection at PC105 was sufficient to enable postnatal identification of injected animals. Lower volumes may also be sufficient but have not been empirically tested.

• Fetal CSF injection of viral vectors may result in transduction of the peripheral nervous system. In such cases, peripheral optogenetic stimulation (e.g., of the tail) in animals expressing opsins may evoke a motor response and/or vocalization, which can aid in identifying transduced animals.

### Problem 7

Uneven or weak viral transduction of the fetus (refer to step 39).

### Potential solution

• Uneven vector distribution following fetal CSF injection may result in preferential transduction of one hemisphere, regional differences in expression, or transduction outside the intended brain regions. When possible, prioritize midline injections when feasible to promote more uniform viral diffusion.

• Deliver the vector incrementally rather than as a single bolus and withdraw the needle slowly and steadily. To minimize backflow, maintain the needle in position for up to 5 minutes before withdrawal and retract the needle slowly and steadily. The minimum dwell time required to prevent backflow has not been systematically evaluated.

• Confirm that injection volume and viral titer are appropriate for the gestational stage. Insufficient volume or low titer may result in reduced or sparse transduction.

• Variability can be managed by screening animals postnatally. When expression levels are sufficiently robust at birth, uneven transduction patterns can be qualitatively assessed using a handheld light source, and animals may be assigned to experiments accordingly.

## Resource availability

### Lead contact

Further information and requests for resources and reagents should be directed to and will be fulfilled by the lead contact, David A. Leopold.

### Technical contact

Further information on executing this protocol should be directed to and will be answered by the technical contact, Ana Rita Ribeiro Gomes.

### Materials availability

Materials used in this protocol are listed in the Key Resources table.

#### Data

All data reported in this paper will be shared by the lead contact upon request.

### Code

This paper does not report original code.

## Additional information

Any additional information is available from the lead contact upon request.

## Acknowledgments

This work was supported by funding from the Intramural Research Program of the National Institute of Mental Health (ZIAMH002898) to D.A.L. and the National Institute of Child Health and Human Development (P50HD103536), University of Rochester Intellectual and Developmental Disabilities Research Center (UR-IDDRC), to K.H.W. A.R.R.G. was funded by the National Institutes of Health’s Visiting Fellow Intramural Research Training Award. N.H. was funded by the National Institutes of Health postbaccalaureate Intramural Research Training Award. We thank George Dold and members of the Section on Instrumentation of the Intramural Research Program of the National Institute of Mental Health for their support in designing and fabricating the custom-made parts used in this project. We are grateful for support from the Systems Neuroscience Imaging Resource of the Intramural Research Program of the National Institute of Mental Health for their allocation of imaging resources used in this research. We also thank the Veterinary Medicine and Resources Branch from the Intramural Research Program of the National Institute of Mental Health for their much-valued support with veterinary health care, animal husbandry, and technical assistance. Finally, we are grateful to Lenegereshe Baweke, Sean Kearney and Naim Wright for technical assistance. This research was supported in part by the Intramural Research Program of the National Institutes of Health (NIH). The contributions of the NIH author(s) are considered Works of the United States Government. The findings and conclusions presented in this paper are those of the authors and do not necessarily reflect the views of the NIH or the U.S. Department of Health and Human Services.

## Author contributions

D.A.L., K.H.W., and A.R.R.G. conceptualized the project. All authors contributed to the development the setup and protocol. A.R.R.G., N.H. and S.M. conducted the procedures. A.R.R.G. collected and processed the data. D.A.L. and K.H.W. acquired funding. D.A.L. and K.H.W. supervised the project.

A.R.R.G. wrote the original draft of the paper. A.R.R.G, D.A.L., and K.H.W. contributed to reviewing and editing the manuscript. All authors gave approval of the final version of the manuscript.

## Declaration of interests

The authors declare no competing interests.

## Methods Video legends

**Methods Video 1: Guide needle sheathing by shrinking Palladium Pebax tubing with a heat gun.** The video shows angled positioning of the guide needle with hemostatic forceps and progressive heat application from the beveled to the blunt end of the needle. Tubing contraction and proximal retraction are visible throughout the process and become more apparent during final contraction at the blunt end.

**Methods Video 2. Ultrasound imaging of a marmoset fetal head acquired using the Vevo MD ultrasound system with the UHF22 transducer.** Scale: 3 mm.

**Methods Video 3. Ultrasound imaging of the same fetus shown in Methods Video 2, acquired using the Clarius L20 HD3 handheld ultrasound system.** Scale: 3 mm.

**Methods Video 4.** Skin penetration facilitated by current pulses applied to the blunt end of the guide tube. Note the transient reduction in echogenicity beneath the guide tube caused by air bubble formation during electrocautery. Ultrasound visualization can be improved by displacing or removing the bubbles. Scale: 1 mm.

**Methods Video 5. Ultrasound-guided FIVI targeting the midline third ventricle in a PC107 marmoset fetus.** The injection trajectory traverses the placenta before entry into the fetal brain through a weaker region of the developing skull. Scale: 1 mm.

**Methods Video 6. Displacement of the fetal head during attempted skull penetration at PC80.** During retraction of the needle, the fetus was repositioned by moving the dam with the translation stages, thereby shifting the fetus toward the left. Scale: 1 mm.

## Supplementary Information

**Data S1. Drawings of the guide tube holder and cradle.**

## Notes

### Competing Interest Statement

The authors have declared no competing interest.

### Summary of Updates

Updated the manuscript to improve its overall organization and readability, including the addition of new sections and reorganization of troubleshooting guidance, and added further information to help users implement the protocol.

